# Obsessive-Compulsive Disorder (OCD) is Associated with Increased Engagement of Frontal Brain Regions Across Multiple Event Related Potentials

**DOI:** 10.1101/2022.11.05.515279

**Authors:** M. Prabhavi N. Perera, Sudaraka Mallawaarachchi, Neil W. Bailey, Oscar W. Murphy, Paul B. Fitzgerald

## Abstract

**Background:** Obsessive-Compulsive Disorder (OCD) is a psychiatric condition leading to significant distress and poor quality of life. Successful treatment of OCD is restricted by the limited knowledge about its pathophysiology. This study aimed to investigate the pathophysiology of OCD using electroencephalographic (EEG) event related potentials (ERP), elicited from multiple tasks to characterise disorder-related differences in underlying brain activity across multiple neural processes.

**Methods:** ERP data were obtained from 25 OCD patients and 27 age- and sex-matched healthy controls (HC) by recording EEG during Flanker and Go/Nogo tasks. Error-related negativity (ERN) was elicited by the Flanker task, while N200 and P300 were generated using the Go/Nogo task. Primary comparisons of the neural response amplitudes and the topographical distribution of neural activity were conducted using scalp field differences across all time points and electrodes.

**Results:** Compared to HC, the OCD group showed altered ERP distributions. Contrasting with the previous literature on ERN and N200 topographies in OCD where fronto-central negative voltages were reported, we detected positive voltages. Additionally, the P300 was found to be less negative in the frontal regions. None of these ERP findings were associated with OCD symptom severity.

**Conclusions:** These results indicate that individuals with OCD show altered frontal neural activity across multiple executive function related processes, supporting the frontal dysfunction theory of OCD. Furthermore, due to the lack of association between altered ERPs and OCD symptom severity, they may be considered potential candidate endophenotypes for OCD.

## 1. INTRODUCTION

Obsessive-compulsive disorder (OCD) is a mental health condition with a lifetime prevalence of 2-3% that causes significant impact on the quality of life of sufferers [1, 2]. OCD is characterised by recurrent, intrusive thoughts (obsessions), often accompanied by repetitive behaviours or mental rituals (compulsions) [3]. The pathophysiology of OCD is poorly understood to date, which has led to poor response to many of the first line treatments [4], and significantly limits the development of more effective novel treatments. Therefore, further research to identify the underlying pathophysiological basis of OCD is crucial.

Electroencephalography (EEG) is an affordable and effective tool to explore the electrophysiology of the brain, and many studies have discovered differences in EEG measures in OCD groups when compared to healthy controls (HC) [5]. Event related potentials (ERP) are voltage changes detected in the EEG that occur as a result of the brain’s time locked response to a stimulus [6]. Several ERPs are known to be altered in OCD groups compared to HC.

The error related negativity (ERN) is conventionally defined as a negative deflection of the EEG that occurs approximately 100-150 ms following an erroneous response [7] and is most commonly measured with executive function and inhibition tasks such as the Eriksen Flanker task [8]. Studies have reported that the ERN amplitude is significantly greater in OCD groups when compared to HC [9-11]. In fact, several studies have proposed ERN as a potential candidate endophenotype for OCD as the ERN enhancement was uncorrelated to symptom severity, and no changes were noted with successful treatment, [12, 13]. However, at least one study has reported no significant difference in the ERN amplitude between OCD and HC groups [14].

Additionally, the ERN is thought to reflect performance and conflict monitoring, where clashes between multiple simultaneously active response tendencies exist, rather than simply a response to having made an error [15]. In fact, previous research has found that the ERN activity starts slightly before the response [16]. Hyperactive behavioural aspects of OCD, including the feeling of incompleteness, doubt and repetitive behaviours may reflect overactive performance monitoring [17]. Furthermore, neuroimaging studies have identified that OCD patients exhibit excessive activity in brain regions that are thought to be associated with performance monitoring, such as the anterior cingulate cortex, orbitofrontal cortex and the dorsolateral prefrontal cortex (DLPFC) [18, 19].

Two other ERPs that are commonly associated with OCD are the N200 and P300, which are both elicited using response inhibition tasks, such as the Go/Nogo task. The N200 is thought to be an ERP that signals the necessity to increase cognitive control to avoid erroneous responses [15], and the P300 is thought to be involved in focusing attention when performing a broad range of cognitive processes [20]. Significantly enhanced N200 amplitudes have been reported in OCD [21, 22]. Compared to HC, the P300 results have been inconsistent with reports of both higher [23, 24] and lower [25, 26] amplitudes in OCD, although overfocused attention has been reported [27].

In the context of these inconsistencies, it is worth noting that previous research on ERP differences in OCD have focused on single electrode analyses and have used inconsistent time windows and tasks [5]. This may have resulted in both inflated false positives and in an inability to detect significant results in alternative time windows [28]. Single electrode analyses are also unable to differentiate between actual neural activity in the observed region, and apparent signal generated by interference patterns between interacting brain regions. This could be theoretically important, as characterising OCD as showing “overactive” error monitoring is based on the perspective that OCD shows enhanced reactions to errors (e.g., larger ERNs), while an altered distribution of activity would suggest qualitative differences in error processing. This difference could have treatment implications, with an increased overall response suggesting treatments to reduce error processing reactions would be valuable, while an altered distribution of activity would suggest treatments that target specific brain regions could be beneficial, and psychological treatments might be optimised by addressing qualitative aspects of error processing rather than the size of the reaction to the errors.

The current study was designed to address the shortcomings of the previous literature by analysing ERP data using an assumption-free technique that encompasses all electrodes and time windows to [29]. The primary aim of the current study was to investigate whether individuals with OCD showed differences in neural activity related to conflict monitoring and inhibitory control when compared to HC. The primary hypothesis was that, compared to HC, the OCD group will show neural alterations in the ERN, N200 and P300 time-windows. Additionally, exploratory analyses were performed to assess for altered distribution of ERP activity in OCD, differences in the P300 amplitude and the relationship between these ERPs and symptom severity. To our knowledge, this is the first study to use analysis techniques that separately tested the overall amplitude differences of neural responses and the topographical distribution of neural activity, while incorporating all electrodes and time windows without a priori assumptions.

## 2. METHODS

### 2.1. Participants

Male and female participants aged between 18 and 65 years were recruited from the state of Victoria, Australia. Written and verbal descriptions of the procedures involved were conveyed to the participants prior to obtaining informed written consent. All participants received a reimbursement for participation. The trial received ethics approval from the Monash Health Human Research Ethics Committee and was conducted under the Good Clinical Practice guidelines [30]. The study was registered in the Australian New Zealand Clinical Trials Registry (ANZCTR; Trial ID: ACTRN12620000748910).

Twenty five Individuals with an OCD diagnosis according to the International Classification of Diseases – 10^th^ revision [31] or DSM-IV/V [3] were included in the OCD group. The Yale Brown Obsessive Compulsive Scale (YBOCS) [32] was used to assess symptom severity, while the Beck Anxiety Inventory (BAI) [33] and the Quick Inventory for Depressive Symptoms – Self Report (QIDS-SR) [34] were used to assess the level of anxiety and concurrent depression, respectively. Exclusion criteria for the OCD group were: (1) presence of an unstable medical/neurological disorder; (2) being diagnosed with another psychiatric condition other than depression/anxiety and (3) scoring <17 on the YBOCS. Participants were recruited regardless of their medication status but were required to be on a stable dose for at least 6 weeks prior to the EEG session. Clinical data recorded from the OCD group included age of onset, duration of illness, presence of comorbidities and medication history.

The HC group included 27 individuals who have never been diagnosed with a psychiatric or neurological illness. HC participants who were currently on psychoactive medications or consuming >2 standard drinks of alcohol per day were excluded.

### 2.2. Tasks and Stimuli

Participants performed two tasks while EEG was recorded: the Go/Nogo task and the Flanker task (**Figure 1**). Stimuli were presented using Inquisit [35] and on a computer screen situated 75-85 cm from the participants’ eyes. All participants were administered a short practice session with 20 trials before performing each task.

**Figure 1.**
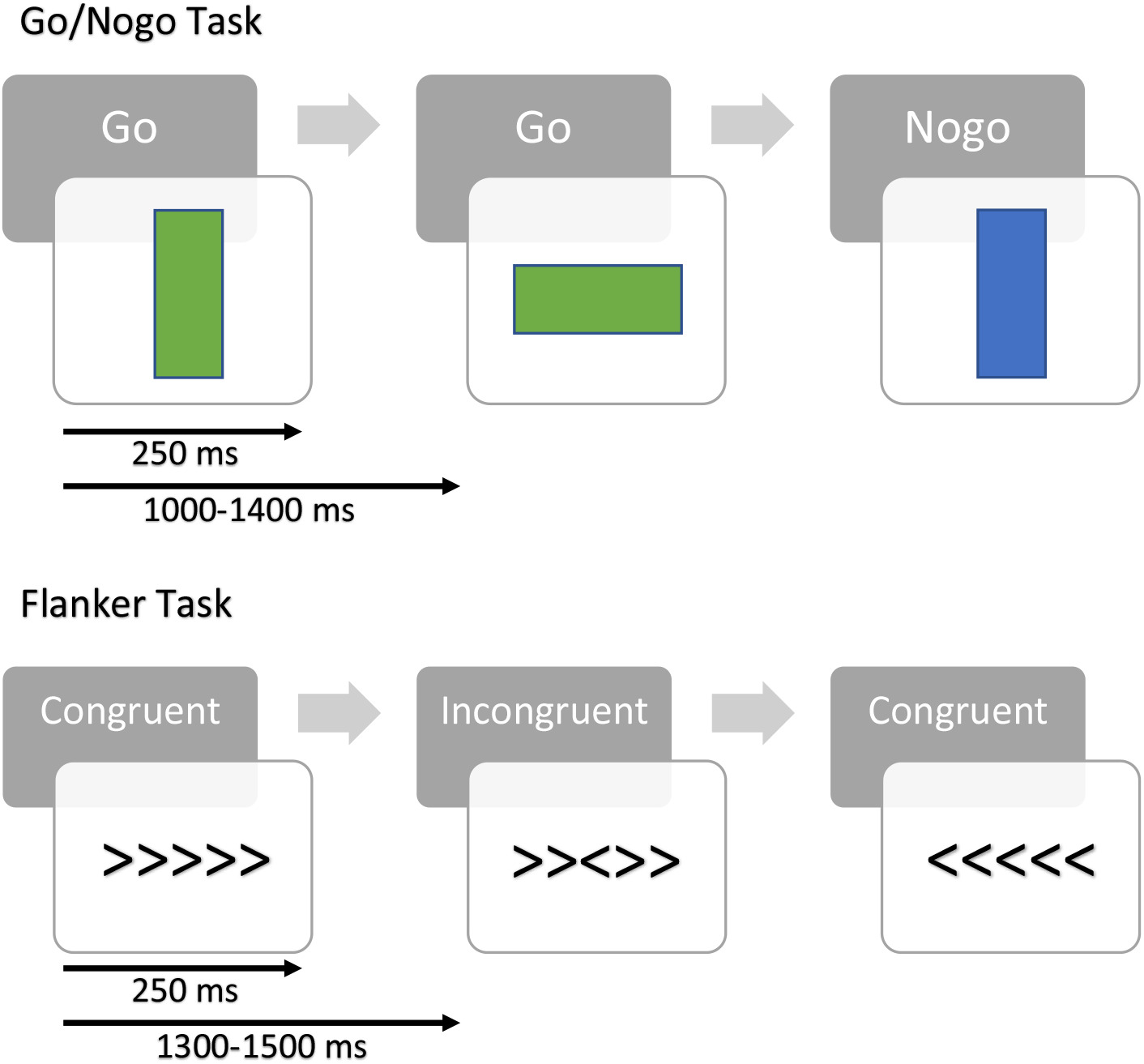
Go/Nogo and Flanker task designs. *Note*. All participants performed two blocks of both tasks, each consisting of 250 trials. Stimuli were presented for 250 ms and the intertrial interval for Go/Nogo and Flanker tasks were 1000-1400 ms and 1300-1500 ms, respectively.

In the Go/Nogo task, participants were requested to respond (Go) to the green rectangle and withhold (Nogo) to the blue. For Go trials, participants were instructed to press the green button with the index finger of the dominant hand as fast as possible. The task included two blocks, each with 250 trials and a break of 1 minute between blocks to avoid fatigue (12 minutes in total). Out of the 250 trials in each block, 20% were Nogo. In each trial, the stimulus was presented for 250 ms with an intertrial interval randomly varying between 1000 and 1400 ms.

In the Flanker task, participants were presented a row of 5 arrows, the middle arrow being the target and the surrounding arrows, flankers. Congruent trials had all five arrows facing the same direction, while incongruent trials had the target arrow facing the opposite direction of the flankers. Participants were instructed to press the right or left button to indicate the direction of the target arrow, while ignoring the flankers. Instructions were also given to respond as quickly and accurately as possible. This task also consisted of two blocks, each with 250 trials and a break of 1 minute between the blocks (12 minutes in total). In each trial, there were equal numbers of left and right targets and equal numbers of congruent and incongruent trials. In each trial, the stimulus was presented for 250 ms with an intertrial interval randomly varying between 1300 and 1500 ms.

### 2.3. EEG recording and Pre-processing

EEG recording was conducted in a laboratory with constant levels of lighting and background noise from air conditioning. Prior to recording, participants were instructed to minimise eye and muscle movements that may affect the EEG recording. Participants were seated upright on a comfortable, padded chair and were requested to stay relaxed during the recording.

EEG was recorded using an actiCHamp amplifier (Brain Products GmbH, Munich, Germany) with BrainVision (version 1.21.0303). The EEG cap included 64 Ag/AgCl electrodes embedded within an EasyCap (Herrsching, Germany) based on the international 10-20 system, out of which 63 electrodes were used for analysis (reference electrode – CPz, ground - AFz). The sampling rate was set at 1000 Hz. A transparent electro-gel was applied onto the scalp at the electrode sites to reduce impedance, which was maintained below 5 kΩ. No online band-pass or notch filtering was applied during the recording.

The recorded, continuous EEG data were pre-processed using the automated RELAX pipeline [36], which was implemented on MATLAB [37] and utilised functions from EEGLAB [38] and fieldtrip [39]. Initially, a 4^th^ order acausal Butterworth bandpass filter from 0.25 to 80 Hz and a 2^nd^ order acausal Butterworth notch filter from 47 to 53 Hz were applied. Subsequently, several measures were taken to detect and reject bad electrodes. The “findNoisyChannels” function of the PREP pipeline was utilised for preliminary removal of noisy channels [40]. Thereafter, marking of electrodes for rejection occurred based on 1) excessive muscle activity [41]; 2) extreme kurtosis; 3) extreme drift; 4) extremely improbable voltage distributions; and 5) extreme outlying amplitudes [42]. Rejection of a maximum of 20% of electrodes was allowed and if >20% were marked for removal, only the worst 20% were removed. The same extreme artifact identification criteria were used after extremely bad electrodes were removed to also mark extreme outlying periods for exclusion from further analysis. Thereafter, three types of artifacts were addressed using Multi-channel Wiener filter (MWF) [43] steps: 1) Muscle activity: epochs affected by muscle activity were identified by low-power log-frequency slopes of >- 0.59; 2) Blink artifacts: pre-specified blink affected channels were selected and voltages were averaged within 1 s epochs after bandpass filtering using a 4^th^ order Butterworth filter from 1 to 25 Hz. Time points where the averaged voltage exceeded the upper quartile from all data, plus thrice the inter-quartile range of all voltages were flagged as blinks. An 800 ms window surrounding each blink was marked as an artifact for cleaning with the MWF; 3) Horizontal eye movements and drift: horizontal eye movements were classified as periods where selected lateral electrodes showed voltages greater than twice the median absolute deviation (MAD) from the median of their overall amplitude, with the same criteria but applied for the opposite voltage polarity in the electrodes on the opposite side of the head [44]. EEG data were considered to be affected by drift if the amplitude was > 10 times MAD from the median of all electrodes [45].

After the MWF cleaning, data were average re-referenced using the PREP method [40], and then subjected to Independent Component Analysis (ICA) using fastICA [46]. Artifactual ICA components were detected using ICLabel [47] and these were cleaned with wavelet enhanced ICA (wICA) [48]. Continuous data were then reconstructed into the scalp space and rejected electrodes were spherically interpolated to obtain a full set of electrodes for all participants. The data were baseline corrected to the -400 to -100 ms period for the Flanker task and from -100 to 0 ms for the Go/Nogo task using the regression baseline correction method [49], applied using an algorithm within the RELAX pipeline [42]. Data were epoched based on the task: 1) for Go/Nogo task, -100 to 500 ms surrounding the onset of the stimulus; 2) for Flanker task, -200 to 400ms surrounding the onset of incorrect responses. Epochs were rejected if the max-min voltage >60 *μ*V or kurtosis/improbable data occurred >3 overall or >5 at any electrode. Each participant was required to have at least 30 epochs in the Go/Nogo task and 6 epochs with errors in the Flanker task to be included in the ERP analysis. One HC and one OCD participant were excluded from the Flanker analysis due to the available epochs with errors being <6. The final sample size for the Flanker analysis was 50 (24 OCD and 26 HC), while all participants were included in the Go/Nogo analysis.

### 2.4. Statistical Comparisons

Behavioural and self-report data were compared using robust tests [50, 51]. An independent sample t-test was used to compare between-group ages and chi-squared tests were used to compare gender, handedness, and marital status. Between-group behavioural performance measures of percentage correct and reaction times were compared using t-tests.

### 2.5. Primary Analysis

The primary statistical comparisons of the ERP data were conducted using the Randomised Graphical User Interface (RAGU), which compares scalp field differences across all time points and electrodes using randomisation statistics [29]. This tool allows comparison of scalp field differences using powerful, assumption-free randomisation statistics between groups and conditions. RAGU controls for experiment-wise multiple comparisons by computing a global duration threshold, which is the 95^th^ percentile of null significant effects, and only real effects with durations longer than this threshold are deemed significant.

Task related data were first imported to RAGU and between- and within-group designs were defined. For both tasks, independent comparisons of the overall strength of scalp field differences and the distribution of neural activity were computed using the global field power (GFP) test and the topographical analysis of variance (TANOVA), respectively. Prior to conducting TANOVA, a topographical consistency test (TCT) was carried out to confirm that scalp activity was distributed consistently within each group and condition. The TCT checks for patterns of consistency in the active sources between the subjects of each tested group and condition [29]. In regions with non-significant TCT results, potential between-group differences can be due to a lack of consistent activation in one or both groups, rather than due to genuine differences. Therefore, significant TCT results provide validity to between-group analyses.

GFP and TANOVA tests were conducted for the Go/Nogo task data as a 2 group (OCD and HC) x 2 condition (Go and Nogo) comparison and as a between-group analysis for the Flanker task data. The number of iterations was set at 5000 with an alpha of *p*= 0.05. The global count p-value examines the likelihood of the overall count of significant time points (at alpha = 0.05) being observed by chance. The global count p-values of all primary and exploratory tests were corrected for experiment-wise multiple comparisons using the Benjamini and Hochberg false discovery rate (FDR) method [52]. Overall p-values for significant periods that passed the global duration threshold were obtained by averaging the p-values of individual time points of that region. Furthermore, p-values were computed averaged across the typical ERN (100 – 150 ms following an erroneous response), N200 (180 – 230 ms post-stimulus) and P300 (250 – 400 ms post-stimulus) windows as defined in the previous literature [7, 53, 54].

To compare with the previous research, average ERP waveforms from the FCz and Pz electrodes were extracted for the ERN/N200 and P300, respectively. Using conventional definitions for each of these ERPs, comparisons were performed using t-tests (supplementary material S1).

### 2.6. TANCOVA

Using data from the OCD group, exploratory analyses were performed to assess the relationship between OCD symptom severity and neural activity findings. The identified significant periods from the TANOVA analysis were averaged and compared using topographical analysis of covariance (TANCOVA).

## 3. RESULTS

### 3.1. Demographic and Behavioural Data

No significant differences were observed in demographic and clinical data between the OCD and HC groups (**Table 1**). The OCD group had a significantly slower reaction time in the Flanker task (*p*= 0.024), but no other significant differences were present in behavioural comparisons (all *p*> 0.05).

**Table 1.** Demographic, Clinical and Behavioural Data of Participants

| Variable | OCD (n = 25) |  |  | HC (n = 27) |  |  | Test statistic (p-val) |
| --- | --- | --- | --- | --- | --- | --- | --- |
|  | Mean | SD | n | Mean | SD | n |  |
| Demographic Data |  |  |  |  |  |  |  |
| Age (years) | 36.24 | 13.06 | 25 | 31.22 | 10.66 | 27 | $t = 1.52, p = 0.13$ |
| Gender (M/F) | | | 12/13 | | | 9/18 | $\chi^2 = 1.14, p = 0.29$ |
| Marital status (S/M) | | | 20/5 | | | 21/6 | $\chi^2 = 0.04, p = 0.85$ |
| Handedness (R/L) | | | 21/4 | | | 26/1 | $\chi^2 = 2.22, p = 0.14$ |
| Clinical Data |  |  |  |  |  |  |  |
| Age at onset (years) | 24.64 | 9.90 | 25 |  |  |  |  |
| Duration of illness (years) | 11.60 | 9.98 | 25 |  |  |  |  |
| YBOCS (total) | 28.00 | 3.82 | 25 |  |  |  |  |
| YBOCS - obsessions | 13.88 | 1.81 | 25 |  |  |  |  |
| YBOCS - compulsions | 14.12 | 2.39 | 25 |  |  |  |  |
| BAI | 17.04 | 8.92 | 24<sup>†</sup> |  |  |  |  |
| QIDS-SR | 10.17 | 4.94 | 24<sup>†</sup> |  |  |  |  |
| Behavioural Data – Flanker Task |  |  |  |  |  |  |  |
| Total percentage correct (%) | 85.39 | 12.76 | 25 | 87.55 | 6.53 | 27 | $t = -0.07, p = 0.965$ |
| Total RT (ms) | 379.38 | 71.12 | 25 | 338.74 | 53.98 | 27 | $t = -2.00, p = 0.045^*$ |
| Correct response RT (ms) | 368.64 | 69.54 | 25 | 339.39 | 49.99 | 27 | $t = -1.33, p = 0.175$ |
| Incorrect response RT (ms) | 376.36 | 124.60 | 25 | 336.17 | 89.87 | 27 | $t = -1.09, p = 0.289$ |
| Behavioural Data – Go/Nogo Task |  |  |  |  |  |  |  |
| Total percentage correct | 90.94 | 7.91 | 25 | 92.64 | 4.65 | 27 | $t = 1.28, p = 0.205$ |
| Go trials RT (ms) | 269.83 | 43.52 | 25 | 261.50 | 42.44 | 27 | $t = 0.09, p = 0.938$ |

OCD – Obsessive Compulsive Disorder, HC – Healthy Control, SD – Standard Deviation, M – Male, F – Female, R – Right, L – Left, S – Single, M – Married, YBOCS – Yale-Brown Obsessive Compulsive Scale, BAI – Beck Anxiety Inventory, QIDS-SR – Quick Inventory of Depressive Symptoms-Self Report, RT = Reaction Time
<sup>†</sup> BAI and QIDS-SR scores for one participant were unavailable due to a data collection issue.
\* Significance level ( $p < 0.05$ )

### 3.2. Topographical Consistency Test

**Figure 2** shows the TCT results for Flanker and Go/Nogo tasks. In the Flanker task, the TCT showed overall signal consistency, except prior to the response and a brief period from 100 to 120 ms in the OCD group. Similarly, in the Go/Nogo task, two brief periods from 153-157 ms and 211-220 ms in the Nogo condition of the OCD group showed evidence of inconsistency. The period lacking consistency in the Flanker task and the latter period of the Nogo condition briefly overlap the significant periods identified as the ERN and N200, which might indicate the between-group difference is due to a lack of consistent signal variability in the OCD group rather than a consistent difference between groups.

**Figure 2.**
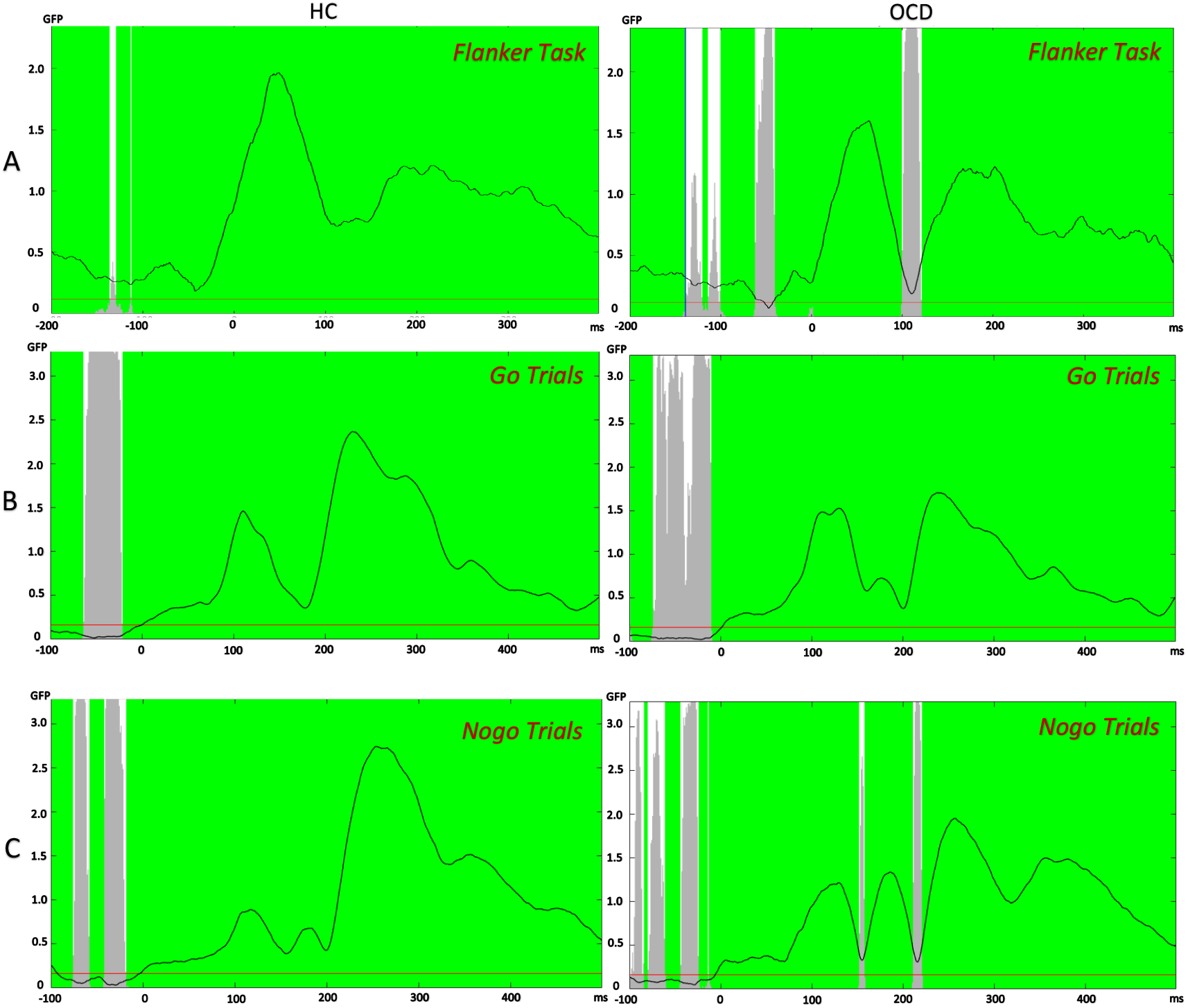
Topographical Consistency Test outcomes for all groups and conditions. *Note*. A) TCT outcome of both OCD and HC groups during the Flanker task. The OCD group showed a brief period with lack of consistency from 100 ms to 120 ms post-response. B) TCT outcome of the Go/Nogo task during Go trials: there was consistency in the signal throughout, except prior to the stimuli. C) TCT outcome of the Go/Nogo task during Nogo trials: there were two brief periods of deficient consistency from 153 ms to 157 ms and 211 ms to 220 ms. Some of these periods overlap the significant period of the ERN and N200, which might reflect a lack of consistent variability in the OCD group rather than an actual consistent difference between groups. (GFP – Global Field Potential, HC – Healthy Control, OCD – Obsessive Compulsive Disorder, TCT – Topographical Consistency Test)

### 3.3. TANOVA

For the Flanker task, main effects of group showed two regions of significance that survived the duration control for multiple comparisons: 1) from -25 to 19 ms (averaged *p*= 0.0071); 2) from 102 to 151 ms, i.e., ERN window (averaged *p*= 0.0178). The global count p = 0.0324 (FDR corrected *p*= 0.032) and the global duration threshold was 38 ms. When the p-values were averaged over the typical ERN window (100 to 150 ms), the difference remained significant (*p*= 0.019). **Figure 3** shows the topographical differences between groups for these two significant windows. Overall, when comparing the ERN window between the HC and OCD groups, the OCD group displays greater frontal positivity (maximal at F3), as well as greater negativity in centroparietal electrodes (maximal at CP4).

**Figure 3.**
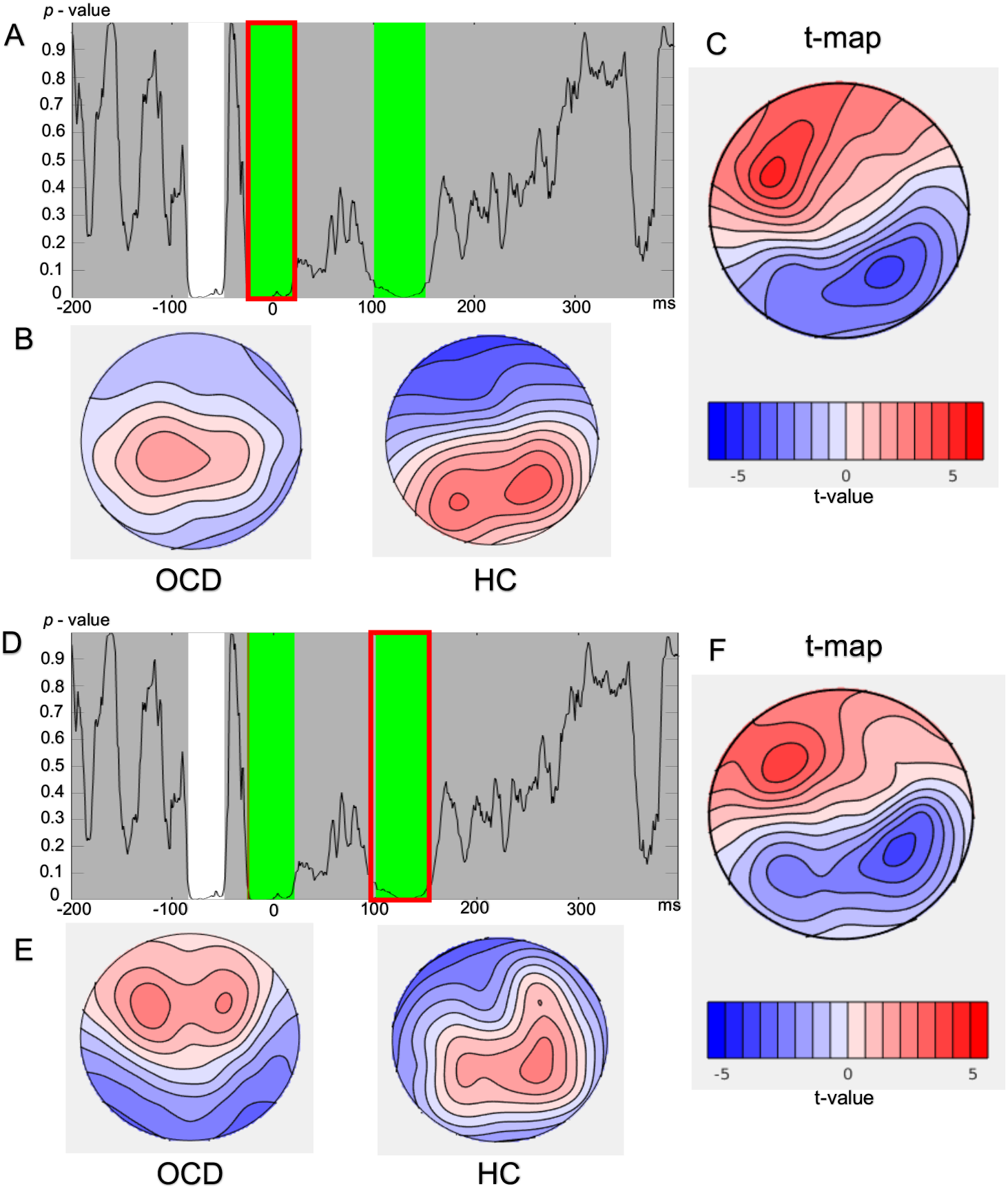
TANOVA main group effect with the flanker task. *Note*. A,D – p values of the between group comparison across the entire epoch of the flanker task. The green highlighted areas (A: -25ms to 19ms, D: 102ms to 151ms) reflect periods that exceed the duration control (38ms) for multiple comparisons across time. B,E – Averaged topographical maps for each group during the significant window. C,F – t-map for topography of the OCD group minus healthy control topography during the significant time window. (OCD – Obsessive Compulsive Disorder, HC – Healthy Control, TANOVA – Topographical Analysis of Variance)

In the Go/Nogo task, the group main effects showed two significantly different regions that passed the duration control for multiple comparisons: 1) from 182 to 230 ms during the N200 window (averaged *p*= 0.0184); 2) from 272 to 323 ms during the P300 window (averaged *p*= 0.0206). The global count p = 0.016 (FDR corrected *p*= 0.032) and the global duration threshold was 44 ms. When the p-values were averaged over typical N200 (180 to 230 ms) and P300 (250 to 400 ms) windows, the differences remained significant for the N200 (*p*= 0.02), but not for the P300 (*p*= 0.185). **Figure 4** depicts the topographical differences between groups for these two significant windows. In the N200 window, when compared to the HC group, the OCD group displayed an atypical distribution, with positive frontal voltages (maximal at AF8), while more negative voltages were found posteriorly (maximal at PO7). Within the P300 window the OCD group also showed an atypical distribution, with stronger positive frontal voltages (maximal at F3), and negative posterior voltages (maximal at P6).

**Figure 4.**
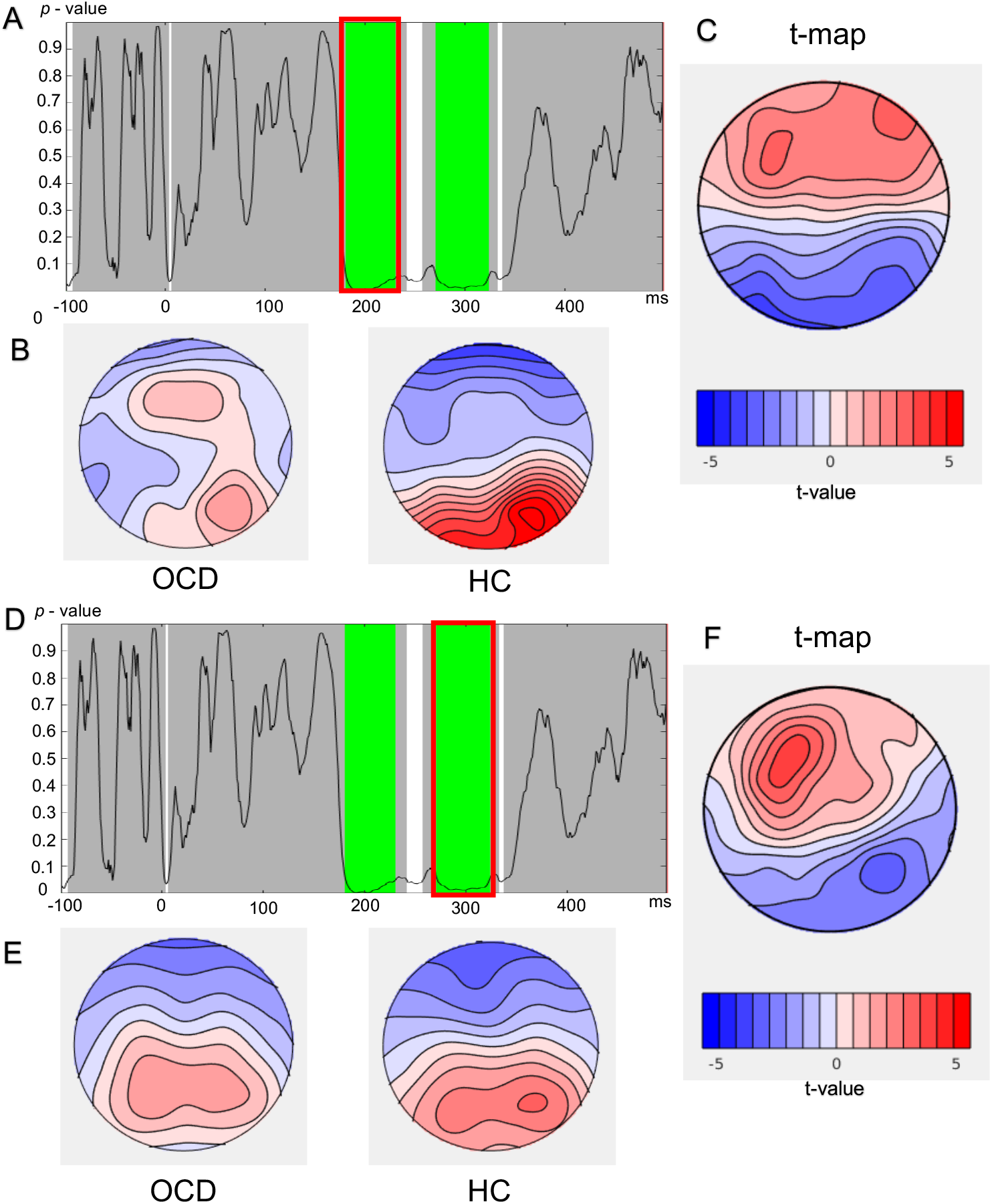
TANOVA main group effect with the Go/Nogo task. *Note*. A,D – p values of the between group comparison across the entire epoch of the Go/Nogo task. The green highlighted areas (A: 182ms to 230ms, D: 272ms to 323ms) reflect periods that exceed the duration control (44ms) for multiple comparisons across time. B,E – Averaged topographical maps for each group during the significant window. C,F – t-map for topography of the OCD group minus healthy control topography during the significant time window. (OCD – Obsessive Compulsive Disorder, HC – Healthy Control, TANOVA – Topographical Analysis of Variance)

### 3.4. Global Field Potential Test

The GFP test was conducted to assess the strength of the neural response to each task. There were no significant time windows that passed the global duration threshold for the Flanker or Go/Nogo tasks for the group main effect or the group (HC, OCD) by condition (Go, Nogo) interaction, indicating that no significant differences were present (all p > 0.05). This result indicates that there were no differences in the overall amplitude of neural responses following errors in the Flanker task or in response to the go/nogo stimuli.

### 3.5. TANCOVA

The TANCOVA between the total YBOCS score and topographical findings across the ERN window showed no significant relationship (*p*= 0.716), indicating that there was no significant association between the ERN and OCD symptom severity. Similarly, no significant relationships were identified between the total YBOCS and the topographical findings of the windows of N200 (*p*= 0.597) or P300 (*p*= 0.281).

## 4. DISCUSSION

The current study examined whether individuals with OCD showed differences in neural activity related to conflict monitoring and response inhibition. The analysis techniques enabled separate examination of differences in the distribution of brain activity and the strength of neural activation, without a priori assumptions about electrodes or time windows. The OCD and HC groups had comparable behavioural performance in both Flanker and Go/Nogo tasks, except for a significantly longer reaction time in the Flanker task in OCD patients. Compared to HC, the OCD group showed more negative voltages centro-parietally for the ERN and posteriorly for the N200 and P300. More positive voltages were noted frontally for all three ERPs. These findings suggest an array of neural differences between OCD and HC groups, which are likely to reflect alterations in executive functions such as attentional processes, conflict monitoring and response inhibition, perhaps produced by frontal dysfunction, which is a known finding in OCD [55, 56].

### 4.1. Error Related Negativity (ERN)

The ERN has been suggested to reflect an error detection and conflict monitoring process, and it provides an evaluation of the consequence of error, which contributes to the adjustment of cognition that serves to prevent future errors [57]. Several previous ERP studies have addressed the question of whether conflict monitoring is enhanced in OCD with reports of significantly elevated ERN [9-11, 58, 59], mostly noted in midline fronto-central electrodes such as FCz, Fz and Cz [14, 60]. In contrast, we found less positive voltages in the ERN around the centroparietal regions, and an alteration to the typical ERN topography such that the OCD group had positive frontal voltages in the ERN window. However, unlike our analyses, these previous single electrode analyses cannot differentiate between differences in the distribution of activity and differences in overall amplitude. As such, our study is the first to show that it is the distribution of the ERN that is altered in OCD rather than the overall amplitude. Moreover, our previous study of spectral power analysis in OCD using the same sample, reported significantly elevated delta and theta power in the same centroparietal electrodes showing higher negativity in the current study [61]. This finding is in agreement with the notion that the ERN emerges, at least in part, from ongoing theta band activity [62, 63], as both seem to be enhanced in OCD when compared to HC.

The conflict hypothesis of the ERN postulates that although ERN may be emitted by the anterior cingulate cortex, response competition processing and greater top-down control processes are recruited from the DLPFC to improve task performance [15, 64]. Additionally, several studies have reported neuroimaging evidence of anomalies in the DLPFC in OCD [19, 65]. Therefore, the altered ERN pattern seen in the current study, with an altered centroparietal and frontotemporal ERN distribution might signify an increased role of the DLPFC in performance monitoring in individuals with OCD.

However, our TCT findings reported an inconsistent period from 100 to 120 ms post-response, which overlaps with the ERN window (102 to 151 ms). This suggests that the significant effect may be at least partly explained by inconsistent neural activation within the OCD group. This has implications for the characterisation of OCD as a disorder with a single, uniquely distinguishable origin common across individuals with OCD, and instead suggests the potential involvement of multiple sources.

The altered distribution of ERN activity in OCD was not found to be related to symptom severity, which supports the theory that an altered ERN may be a candidate endophenotype for OCD. Endophenotypes are defined as objectively measurable elements that are related to an underlying susceptibility for a disease, and are characterised by several factors: endophenotypes, 1) are related to the illness in the population, 2) manifest independent of the presence of symptoms, 3) are heritable, 4) may be present in unaffected relatives and 5) are co-segregated with the illness within families [66]. The first report of ERN as a candidate endophenotype for OCD was in a paediatric study, where successful therapy of OCD did not result in a change in the ERN [12]. Consequently, asymptomatic siblings [67] and first-degree relatives [13] of OCD patients were also noted to have significantly enhanced ERN. Furthermore, several studies [68-72], including the present study reported no association between an altered ERN and OCD symptom severity. Therefore, our findings support the concept that ERN may be a candidate endophenotype for OCD. Our results also suggest that the characterisation of the ERN as an endophenotype is likely to be more sophisticated than can be provided by traditional single electrode analyses.

### 4.2. N200 and P300

The N200 is reported to reflect processes underlying response inhibition and conflict monitoring, both of which are known to be dysfunctional in OCD groups [73-75]. Our findings indicate that in the OCD group, the N200 had a distribution of neural activity that differed from the typical N200 topography, with positive fronto-central voltages and more negative posterior voltages when compared to HC. These results conflict with reports of several studies that found significantly larger N200 amplitudes in OCD groups when compared to HC [21, 22, 76-78]. The frontocentral electrodes that were detected to have a positive voltage in our study were previously reported as negative. However, similar to studies of the ERN, these studies typically only assessed midline frontocentral electrodes in their analyses. Instead, our results are consistent with two studies that reported opposing results with significantly smaller N200 amplitudes [79, 80]. Additionally, our single electrode N200 analysis of the FCz electrode (supplementary material S1) showed no significant difference between groups.

The OCD group was found to show less negative frontal voltages in the P300 compared to HC, which is consistent with findings of several previous studies [23, 24, 81, 82]. An altered P300 has been suggested to point to disruptions in the functionality of the brain systems that are required to provide sustained attention [20, 83], and has been found to underpin clinical symptoms of waning attention in mental health conditions such as schizophrenia [84]. Individuals with OCD are also known to have deficits in cognitive processes of attention and selective attention [85, 86]. Therefore, the observed P300 alterations might be associated with sustained attention deficits in OCD. However, a few studies reported contradicting results of lower P300 amplitudes in OCD [25, 26, 78, 87]. These discrepancies in findings may be due to methodological variations; mainly usage of different tasks (Go/Nogo, auditory oddball, visual novelty recognition tasks) to elicit the ERPs.

### 4.3. Limitations and Future Directions

The findings of the current study should be interpreted taking its limitations into account. While the recruited sample size was sufficient to ascertain group-level differences, inclusion of more participants would likely increase the statistical power of the findings, and perhaps reveal additional findings in different time windows after stimulus presentation. Our study also included participants who were on different classes of medications, which potentially increased the heterogeneity of our sample. Previous studies have reported that medications may cause alterations in ERPs [88], and therefore, the recorded EEG findings might be influenced by medication effects. Alternatively, studies exclusively including drug naïve participants should be performed in the future.

Although the ERP differences reported in the current study are known to reflect alterations in several domains of cognition, such as conflict monitoring, response inhibition and attentional processes, our behavioural performance findings were largely non-significant, with the exception that the OCD group showed slower reaction times in the Flanker task. This may be due to the cognitive tasks not being sensitive enough to provide significant behavioural differences or due to insufficient sample size.

Our findings support the notion that the ERN might be a potential candidate endophenotype for OCD, as the altered ERN distribution was not associated with OCD symptom severity. However, other criteria characterising endophenotypes, such as effects of treatment on ERN and presence of raised ERN in first degree relatives were not tested. Although several studies have assessed these criteria partially [12, 67], a comprehensive study assessing all parameters has not been performed to date. Furthermore, treatment of OCD with non-invasive brain stimulation methods such as repetitive transcranial magnetic stimulation (rTMS) has shown to be efficacious [89]. It has been reported that administering TMS may alter the electrophysiology of the brain leading to various EEG changes [90]. Therefore, in addition to assessing medication effects, future rTMS studies should investigate ERP changes pre- to post-stimulation, to confirm whether ERN is indeed an endophenotype for OCD. Our findings indicated that the N200 enhancement was not associated with OCD symptom severity, which is characteristic of an endophenotype. Future studies could also investigate the N200 as a potential endophenotype.

### 4.4. Conclusions

OCD is a mental health condition leading to significant distress and poor quality of life to sufferers. Successful treatment of OCD is restricted due to the limited knowledge on its pathophysiology. The current study aimed to investigate ERP findings of OCD to understand the differences in neural activity which might underlie the disease. ERP data were obtained from 25 OCD patients and 27 HC by recording EEG during Flanker and Go/Nogo tasks. Primary comparisons were conducted using the RAGU interface, which analysed scalp field differences across all time points and electrodes using randomisation statistics. Compared to HC, OCD group showed differences in the distribution of neural activity in the ERN, N200 and P300 windows. TANOVA results indicated less typical ERN and N200 distributions of activity with more positive voltages in frontal electrodes, and more negative voltages centro-parietally and posteriorly in the OCD group. The P300 was also found to be less negative in the frontotemporal regions in OCD. These ERP findings were not associated with OCD symptom severity. The findings of the current study indicate that individuals with OCD show an altered distribution of neural activity related to conflict monitoring and inhibitory control, but not an alteration to overall neural response strength. Furthermore, due to the lack of association between the altered ERPs and OCD symptom severity, the altered distributions of neural activity may be considered as potential candidate endophenotypes for OCD.

## ACKNOWLEDGMENTS

This work was supported by the Monash University Institute of Graduate Research (MPNP) and National Health and Medical Research Council of Australia Investigator grant 1193596 (PBF).

The authors assert that all procedures contributing to this work comply with the ethical standards of the relevant national and institutional committees on human experimentation and with the Helsinki Declaration of 1975, as revised in 2008. This study was approved by the Monash Health Human Research Ethics Committee (Reference: RES-20-0000-265A).

## AUTHOR CONTRIBUTIONS

MPNP – project design, data collection, analysis, and interpretation, writing the manuscript SM – data analysis, review of the manuscript

NWB – data analysis and interpretation, review of the manuscript OWM – data analysis and interpretation, review of the manuscript PBF – project design, data interpretation, review of the manuscript

## REFERENCES

[1] A. Ruscio, D. Stein, W. Chiu, and R. Kessler, “The epidemiology of obsessive-compulsive disorder in the National Comorbidity Survey Replication,” Molecular psychiatry, vol. 15, no. 1, p. 53, 2010.

[2] L. M. Koran, M. L. Thienemann, and R. Davenport, “Quality of life for patients with obsessive-compulsive disorder,” The American Journal of Psychiatry, vol. 153, no. 6, p. 783, 1996.

[3] A. P. Association, Diagnostic and statistical manual of mental disorders (DSM-5®). American Psychiatric Pub, 2013.

[4] S. Taylor, J. S. Abramowitz, and D. McKay, “Non-adherence and non-response in the treatment of anxiety disorders,” Journal of anxiety disorders, vol. 26, no. 5, pp. 583–589, 2012.

[5] M. P. N. Perera, N. W. Bailey, S. E. Herring, and P. B. Fitzgerald, “Electrophysiology of obsessive compulsive disorder: a systematic review of the electroencephalographic literature,” Journal of anxiety disorders, vol. 62, pp. 1–14, 2019.

[6] M. G. Coles and M. D. Rugg, Event-related brain potentials: An introduction. Oxford University Press, 1995.

[7] B. Stemmer, S. J. Segalowitz, W. Witzke, and P. W. Schönle, “Error detection in patients with lesions to the medial prefrontal cortex: an ERP study,” Neuropsychologia, vol. 42, no. 1, pp. 118–130, 2004.

[8] B. A. Eriksen and C. W. Eriksen, “Effects of noise letters upon the identification of a target letter in a nonsearch task,” Perception & psychophysics, vol. 16, no. 1, pp. 143–149, 1974.

[9] T. Endrass, B. Schuermann, C. Kaufmann, R. Spielberg, R. Kniesche, and N. Kathmann, “Performance monitoring and error significance in patients with obsessive-compulsive disorder,” Biological psychology, vol. 84, no. 2, pp. 257–263, 2010.

[10] Z. Xiao et al., “Error-related negativity abnormalities in generalized anxiety disorder and obsessive–compulsive disorder,” Progress in Neuro-Psychopharmacology and Biological Psychiatry, vol. 35, no. 1, pp. 265–272, 2011.

[11] S. Johannes et al., “Discrepant target detection and action monitoring in obsessive–compulsive disorder,” Psychiatry Research: Neuroimaging, vol. 108, no. 2, pp. 101–110, 2001.

[12] G. Hajcak, M. E. Franklin, E. B. Foa, and R. F. Simons, “Increased error-related brain activity in pediatric obsessive-compulsive disorder before and after treatment,” American Journal of Psychiatry, vol. 165, no. 1, pp. 116–123, 2008.

[13] A. Riesel, T. Endrass, C. Kaufmann, and N. Kathmann, “Overactive error-related brain activity as a candidate endophenotype for obsessive-compulsive disorder: evidence from unaffected first-degree relatives,” American Journal of Psychiatry, vol. 168, no. 3, pp. 317–324, 2011.

[14] S. Nieuwenhuis, M. M. Nielen, N. Mol, G. Hajcak, and D. J. Veltman, “Performance monitoring in obsessive-compulsive disorder,” Psychiatry Research, vol. 134, no. 2, pp. 111–122, 2005.

[15] M. M. Botvinick, T. S. Braver, D. M. Barch, C. S. Carter, and J. D. Cohen, “Conflict monitoring and cognitive control,” Psychological review, vol. 108, no. 3, p. 624, 2001.

[16] N. Yeung, M. M. Botvinick, and J. D. Cohen, “The neural basis of error detection: conflict monitoring and the error-related negativity,” Psychological review, vol. 111, no. 4, p. 931, 2004.

[17] R. K. Pitman, “A cybernetic model of obsessive-compulsive psychopathology,” Comprehensive psychiatry, vol. 28, no. 4, pp. 334–343, 1987.

[18] C. S. Carter, T. S. Braver, D. M. Barch, M. M. Botvinick, D. Noll, and J. D. Cohen, “Anterior cingulate cortex, error detection, and the online monitoring of performance,” Science, vol. 280, no. 5364, pp. 747–749, 1998.

[19] A. Russell et al., “Localized functional neurochemical marker abnormalities in dorsolateral prefrontal cortex in pediatric obsessive-compulsive disorder,” Journal of Child and Adolescent Psychopharmacology, vol. 13, no. 2, Supplement 1, pp. 31–38, 2003.

[20] D. E. Linden, “The P300: where in the brain is it produced and what does it tell us?,” The Neuroscientist, vol. 11, no. 6, pp. 563–576, 2005.

[21] A. Riesel, J. Klawohn, N. Kathmann, and T. Endrass, “Conflict monitoring and adaptation as reflected by N2 amplitude in obsessive–compulsive disorder,” Psychological medicine, vol. 47, no. 8, pp. 1379–1388, 2017.

[22] A. Miyata, H. Matsunaga, N. Kiriike, Y. Iwasaki, Y. Takei, and S. Yamagami, “Event-related potentials in patients with obsessive-compulsive disorder,” Psychiatry and clinical neurosciences, vol. 52, no. 5, pp. 513–518, 1998.

[23] C. Andreou et al., “P300 in obsessive–compulsive disorder: source localization and the effects of treatment,” Journal of psychiatric research, vol. 47, no. 12, pp. 1975–1983, 2013.

[24] M. Ischebeck, T. Endrass, D. Simon, and N. Kathmann, “Auditory novelty processing is enhanced in obsessive–compulsive disorder,” Depression and anxiety, vol. 28, no. 10, pp. 915–923, 2011.

[25] M. Sanz, V. Molina, M. Martin-Loeches, A. Calcedo, and F. J. Rubia, “Auditory P300 event related potential and serotonin reuptake inhibitor treatment in obsessive-compulsive disorder patients,” Psychiatry Research, vol. 101, no. 1, pp. 75–81, 2001.

[26] K. Yamamuro et al., “Event-related potentials in drug-naive pediatric patients with obsessive-compulsive disorder,” Psychiatry research, vol. 230, no. 2, pp. 394–399, 2015.

[27] D. J. Stein and N. Fineberg, Obsessive-compulsive disorder. Oxford University Press, 2007.

[28] J. Kilner, “Bias in a common EEG and MEG statistical analysis and how to avoid it,” Clinical Neurophysiology, vol. 10, no. 124, pp. 2062–2063, 2013.

[29] T. Koenig, M. Kottlow, M. Stein, and L. Melie-García, “Ragu: a free tool for the analysis of EEG and MEG event-related scalp field data using global randomization statistics,” Computational intelligence and neuroscience, vol. 2011, 2011.

[30] J. R. Dixon, “The international conference on harmonization good clinical practice guideline,” Quality Assurance, vol. 6, no. 2, pp. 65–74, 1999.

[31] W. H. Organization, The ICD-10 classification of mental and behavioural disorders: diagnostic criteria for research. World Health Organization, 1993.

[32] W. K. Goodman et al., “The yale-brown obsessive compulsive scale: II. Validity,” Archives of general psychiatry, vol. 46, no. 11, pp. 1012–1016, 1989.

[33] A. T. Beck, N. Epstein, G. Brown, and R. A. Steer, “An inventory for measuring clinical anxiety: psychometric properties,” Journal of consulting and clinical psychology, vol. 56, no. 6, p. 893, 1988.

[34] A. J. Rush et al., “The 16-Item Quick Inventory of Depressive Symptomatology (QIDS), clinician rating (QIDS-C), and self-report (QIDS-SR): a psychometric evaluation in patients with chronic major depression,” Biological psychiatry, vol. 54, no. 5, pp. 573–583, 2003.

[35] Millisecond, “Inquisit (Version 4)[Computer software],” 2015.

[36] N. Bailey et al., “Introducing RELAX (the Reduction of Electroencephalographic Artifacts): A fully automated pre-processing pipeline for cleaning EEG data-Part 1: Algorithm and Application to Oscillations,” bioRxiv, 2022.

[37] (2022). The MathWorks Inc., Natick, Massachusetts.

[38] A. Delorme and S. Makeig, “EEGLAB: an open source toolbox for analysis of single-trial EEG dynamics including independent component analysis,” Journal of neuroscience methods, vol. 134, no. 1, pp. 9–21, 2004.

[39] R. Oostenveld, P. Fries, E. Maris, and J.-M. Schoffelen, “FieldTrip: open source software for advanced analysis of MEG, EEG, and invasive electrophysiological data,” Computational intelligence and neuroscience, vol. 2011, 2011.

[40] N. Bigdely-Shamlo, T. Mullen, C. Kothe, K.-M. Su, and K. A. Robbins, “The PREP pipeline: standardized preprocessing for large-scale EEG analysis,” Frontiers in neuroinformatics, vol. 9, p. 16, 2015.

[41] S. Fitzgibbon et al., “Automatic determination of EMG-contaminated components and validation of independent component analysis using EEG during pharmacologic paralysis,” Clinical neurophysiology, vol. 127, no. 3, pp. 1781–1793, 2016.

[42] N. Bailey et al., “Introducing RELAX (the Reduction of Electroencephalographic Artifacts): A fully automated pre-processing pipeline for cleaning EEG data-Part 2: Application to Event-Related Potentials,” bioRxiv, 2022.

[43] B. Somers, T. Francart, and A. Bertrand, “MWF toolbox for EEG artifact removal,” 2019.

[44] N. C. Rogasch et al., “Analysing concurrent transcranial magnetic stimulation and electroencephalographic data: A review and introduction to the open-source TESA software,” Neuroimage, vol. 147, pp. 934–951, 2017.

[45] H. Nolan, R. Whelan, and R. B. Reilly, “FASTER: fully automated statistical thresholding for EEG artifact rejection,” Journal of neuroscience methods, vol. 192, no. 1, pp. 152–162, 2010.

[46] A. Hyvarinen, “Fast ICA for noisy data using Gaussian moments,” in 1999 IEEE international symposium on circuits and systems (ISCAS), 1999, vol. 5: IEEE, pp. 57–61.

[47] L. Pion-Tonachini, K. Kreutz-Delgado, and S. Makeig, “ICLabel: An automated electroencephalographic independent component classifier, dataset, and website,” NeuroImage, vol. 198, pp. 181–197, 2019.

[48] N. P. Castellanos and V. A. Makarov, “Recovering EEG brain signals: Artifact suppression with wavelet enhanced independent component analysis,” Journal of neuroscience methods, vol. 158, no. 2, pp. 300–312, 2006.

[49] P. M. Alday, “How much baseline correction do we need in ERP research? Extended GLM model can replace baseline correction while lifting its limits,” Psychophysiology, vol. 56, no. 12, p. e13451, 2019.

[50] K. K. Yuen, “The two-sample trimmed t for unequal population variances,” Biometrika, vol. 61, no. 1, pp. 165–170, 1974.

[51] P. Mair and R. Wilcox, “Robust statistical methods using WRS2,” The WRS2 Package, 2019.

[52] Y. Benjamini and Y. Hochberg, “Controlling the false discovery rate: a practical and powerful approach to multiple testing,” Journal of the Royal statistical society: series B (Methodological), vol. 57, no. 1, pp. 289–300, 1995.

[53] J. Polich, “On the relationship between EEG and P300: individual differences, aging, and ultradian rhythms,” International journal of psychophysiology, vol. 26, no. 1-3, pp. 299–317, 1997.

[54] A. Brunellière, C. Sánchez-García, N. Ikumi, and S. Soto-Faraco, “Visual information constrains early and late stages of spoken-word recognition in sentence context,” International Journal of Psychophysiology, vol. 89, no. 1, pp. 136–147, 2013.

[55] S. Khanna, “Obsessive-compulsive disorder: Is there a frontal lobe dysfunction?,” Biological Psychiatry, vol. 24, no. 5, pp. 602–613, 1988.

[56] K. Schmidtke, A. Schorb, G. Winkelmann, and F. Hohagen, “Cognitive frontal lobe dysfunction in obsessive-compulsive disorder,” Biological psychiatry, vol. 43, no. 9, pp. 666–673, 1998.

[57] W. J. Gehring, B. Goss, M. G. Coles, D. E. Meyer, and E. Donchin, “A neural system for error detection and compensation,” Psychological science, vol. 4, no. 6, pp. 385–390, 1993.

[58] R. Grützmann, T. Endrass, C. Kaufmann, E. Allen, T. Eichele, and N. Kathmann, “Presupplementary motor area contributes to altered error monitoring in obsessive-compulsive disorder,” Biological psychiatry, vol. 80, no. 7, pp. 562–571, 2016.

[59] D. Roh, J.-G. Chang, S. Yoo, J. Shin, and C.-H. Kim, “Modulation of error monitoring in obsessive–compulsive disorder by individually tailored symptom provocation,” Psychological medicine, vol. 47, no. 12, pp. 2071–2080, 2017.

[60] Z.-M. Zhang et al., “Attentional avoidance of threats in obsessive compulsive disorder: An event related potential study,” Behaviour research and therapy, vol. 97, pp. 96–104, 2017.

[61] M. P. N. Perera, S. Mallawaarachchi, N. W. Bailey, O. W. Murphy, and P. B. Fitzgerald, “Obsessive-Compulsive Disorder (OCD) is Associated with Increased Electroencephalographic (EEG) Delta and Theta Oscillatory Power but Reduced Delta Connectivity,” bioRxiv, p. 2022.10.03.510571, 2022, doi: 10.1101/2022.10.03.510571.

[62] L. T. Trujillo and J. J. Allen, “Theta EEG dynamics of the error-related negativity,” Clinical Neurophysiology, vol. 118, no. 3, pp. 645–668, 2007.

[63] J. F. Cavanagh, L. Zambrano-Vazquez, and J. J. Allen, “Theta lingua franca: A common mid-frontal substrate for action monitoring processes,” Psychophysiology, vol. 49, no. 2, pp. 220–238, 2012.

[64] D. H. Mathalon, S. L. Whitfield, and J. M. Ford, “Anatomy of an error: ERP and fMRI,” Biological psychology, vol. 64, no. 1-2, pp. 119–141, 2003.

[65] S. J. De Wit et al., “Multicenter voxel-based morphometry mega-analysis of structural brain scans in obsessive-compulsive disorder,” American journal of psychiatry, vol. 171, no. 3, pp. 340–349, 2014.

[66] I. I. Gottesman and T. D. Gould, “The endophenotype concept in psychiatry: etymology and strategic intentions,” American Journal of Psychiatry, vol. 160, no. 4, pp. 636–645, 2003.

[67] M. Carrasco, S. M. Harbin, J. K. Nienhuis, K. D. Fitzgerald, W. J. Gehring, and G. L. Hanna, “Increased error-related brain activity in youth with obsessive-compulsive disorder and unaffected siblings,” Depression and Anxiety, vol. 30, no. 1, pp. 39–46, 2013.

[68] T. Endrass, A. Riesel, N. Kathmann, and U. Buhlmann, “Performance monitoring in obsessive–compulsive disorder and social anxiety disorder,” Journal of Abnormal Psychology, vol. 123, no. 4, p. 705, 2014.

[69] G. L. Hanna et al., “Error-related negativity and tic history in pediatric obsessive-compulsive disorder,” Journal of the American Academy of Child & Adolescent Psychiatry, vol. 51, no. 9, pp. 902–910, 2012.

[70] H. Nawani et al., “Enhanced error related negativity amplitude in medication-naïve, comorbidity-free obsessive compulsive disorder,” Psychiatry research, 2017.

[71] A. Riesel, N. Kathmann, and T. Endrass, “Overactive performance monitoring in obsessive–compulsive disorder is independent of symptom expression,” European archives of psychiatry and clinical neuroscience, vol. 264, no. 8, pp. 707–717, 2014.

[72] M. Ruchsow, G. Grön, K. Reuter, M. Spitzer, L. Hermle, and M. Kiefer, “Error-related brain activity in patients with obsessive-compulsive disorder and in healthy controls,” Journal of Psychophysiology, vol. 19, no. 4, pp. 298–304, 2005.

[73] S. R. Chamberlain, A. D. Blackwell, N. A. Fineberg, T. W. Robbins, and B. J. Sahakian, “The neuropsychology of obsessive compulsive disorder: the importance of failures in cognitive and behavioural inhibition as candidate endophenotypic markers,” Neuroscience & Biobehavioral Reviews, vol. 29, no. 3, pp. 399–419, 2005.

[74] J. Woolley, I. Heyman, M. Brammer, I. Frampton, P. K. McGuire, and K. Rubia, “Brain activation in paediatric obsessive-compulsive disorder during tasks of inhibitory control,” The British Journal of Psychiatry, vol. 192, no. 1, pp. 25–31, 2008.

[75] R. Penades, R. Catalan, K. Rubia, S. Andres, M. Salamero, and C. Gasto, “Impaired response inhibition in obsessive compulsive disorder,” European Psychiatry, vol. 22, no. 6, pp. 404–410, 2007.

[76] J. Towey et al., “Endogenous event-related potentials in obsessive-compulsive disorder,” Biological Psychiatry, vol. 28, no. 2, pp. 92–98, 1990.

[77] J. Towey et al., “Event-related potential and clinical correlates of neurodysfunction in obsessive-compulsive disorder,” Psychiatry Research, vol. 49, no. 2, pp. 167–181, 1993.

[78] J. P. Towey et al., “Brain event-related potential correlates of overfocused attention in obsessive-compulsive disorder,” Psychophysiology, vol. 31, no. 6, pp. 535–543, 1994.

[79] P. Morault, F. Guillem, M. Bourgeois, and J. Paty, “Improvement predictors in obsessive-compulsive disorder. An event-related potential study,” Psychiatry research, vol. 81, no. 1, pp. 87–96, 1998.

[80] M. S. Kim, Y. Y. Kim, S. Y. Yoo, and J. S. Kwon, “Electrophysiological correlates of behavioral response inhibition in patients with obsessive–compulsive disorder,” Depression and Anxiety, vol. 24, no. 1, pp. 22–31, 2007.

[81] P. Mavrogiorgou et al., “P300 subcomponents in obsessive-compulsive disorder,” Journal of Psychiatric Research, vol. 36, no. 6, pp. 399–406, 2002.

[82] D. Gohle et al., “Electrophysiological evidence for cortical abnormalities in obsessive–compulsive disorder–A replication study using auditory event-related P300 subcomponents,” Journal of psychiatric research, vol. 42, no. 4, pp. 297–303, 2008.

[83] A. Behzadnia, F. Ghassemi, S. A. Chermahini, Z. Tabanfar, and A. Taymourtash, “The neural correlation of sustained attention in performing conjunctive continuous performance task: an event-related potential study,” Neuroreport, vol. 29, no. 11, pp. 954–961, 2018.

[84] J. M. Ford, “Schizophrenia: the broken P300 and beyond,” Psychophysiology, vol. 36, no. 6, pp. 667–682, 1999.

[85] F. De Geus, D. A. Denys, M. M. Sitskoorn, and H. G. Westenberg, “Attention and cognition in patients with obsessive–compulsive disorder,” Psychiatry and clinical neurosciences, vol. 61, no. 1, pp. 45–53, 2007.

[86] I. C. Clayton, J. C. Richards, and C. J. Edwards, “Selective attention in obsessive–compulsive disorder,” Journal of Abnormal Psychology, vol. 108, no. 1, p. 171, 1999.

[87] P. Malloy, S. Rasmussen, W. Braden, and R. J. Haier, “Topographic evoked potential mapping in obsessive-compulsive disorder: evidence of frontal lobe dysfunction,” Psychiatry Research, vol. 28, no. 1, pp. 63–71, 1989.

[88] J. M. Ford, P. M. White, J. G. Csernansky, W. O. Faustman, W. T. Roth, and A. Pfefferbaum, “ERPs in schizophrenia: effects of antipsychotic medication,” Biological psychiatry, vol. 36, no. 3, pp. 153–170, 1994.

[89] M. P. N. Perera, S. Mallawaarachchi, A. Miljevic, N. W. Bailey, S. E. Herring, and P. B. Fitzgerald, “Repetitive Transcranial Magnetic Stimulation for Obsessive-Compulsive Disorder: A Meta-analysis of Randomized, Sham-Controlled Trials,” Biological Psychiatry: Cognitive Neuroscience and Neuroimaging, vol. 6, no. 10, pp. 947–960, 2021.

[90] G. Thut and A. Pascual-Leone, “A review of combined TMS-EEG studies to characterize lasting effects of repetitive TMS and assess their usefulness in cognitive and clinical neuroscience,” Brain topography, vol. 22, no. 4, pp. 219–232, 2010.

