## Supplementary material S1 for "Obsessive-Compulsive Disorder (OCD) is Associated with Increased Engagement of Frontal Brain Regions Across Multiple Event Related Potentials"

S1. To enable comparisons with results from previous research, the average ERP waveforms for the FCz electrode were also sketched and analysed. Furthermore, the average waveforms were drawn for the electrodes that showed the maximal difference between groups for each task. Independent sample t-tests were conducted to compare groups. For this analysis, ERN was defined as the peak-to-peak difference in voltage between the most positive peak from -100 to 0 ms and the most negative peak from 0 to 150 ms and after an erroneous response in the flanker task [1]. The N200 was defined as the voltage value of the most negative peak between 150 and 250 ms post-stimulus [2] and the P300 as the voltage value of the most positive peak between 250 and 500 ms post-stimulus of the Go/Nogo task [3].

The ERN was found to be significantly more negative in the OCD group when compared to HC at the C6 electrode (t-statistic = 0.3058, p = 0.0115). However, the ERN was not significantly different between groups at FCz (t-statistic = -0.7353, p = 0.2328). The N200 was significantly more negative in the OCD group at the P2 electrode (t-statistic = 3.1614, p = 0.0027), while no differences were noted at FCz (t-statistic = 0.3058, p = 0.7611). The P300 was significantly higher in OCD when compared to HC at the F3 electrode (t-statistic = -2.95, p = 0.0048), while no differences were noted at Pz (t-statistic = -0.21, p = 0.836). Averaged waveforms of each of the above analyses are shown in **Figure S1**.

**Figure S1**

*Single electrode analysis of ERN, N200 and P300*

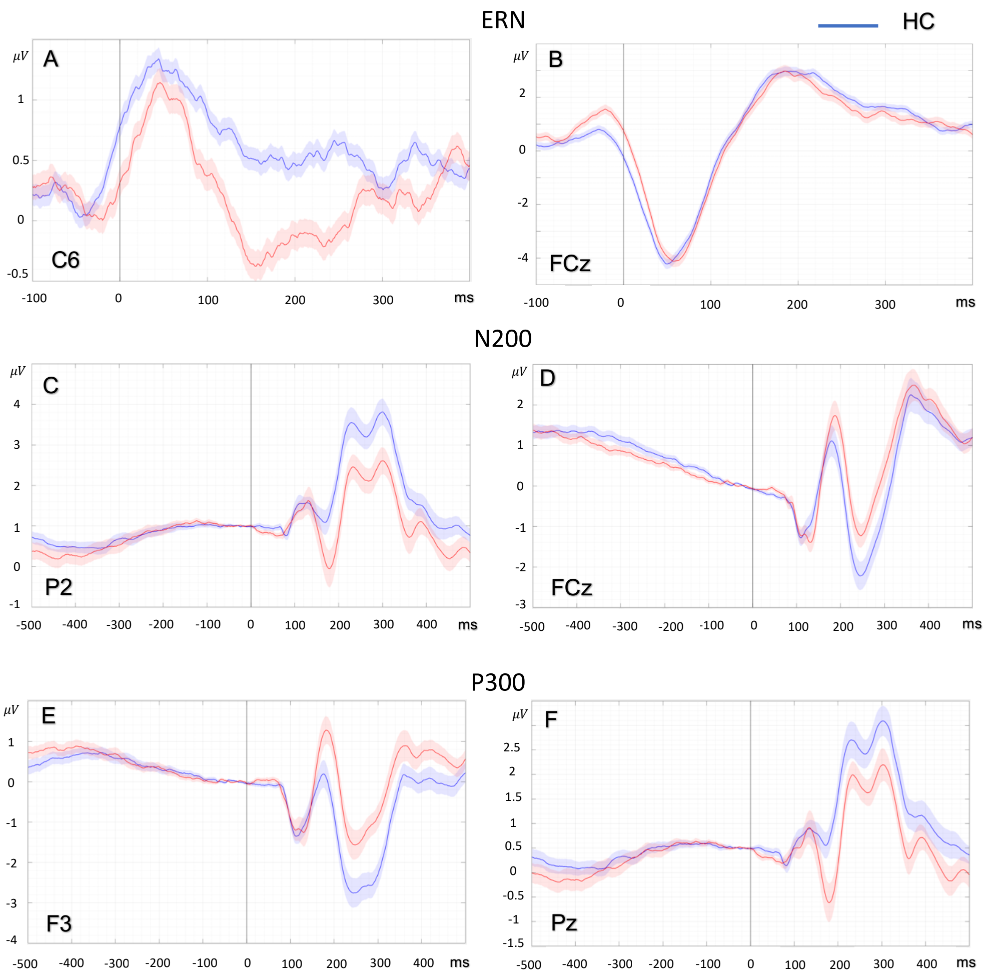


*Note.* Single electrode analyses of ERN, N200 and P300. A) The ERN was found to be significantly more negative in OCD groups than HC at C6 and, B) no between-group difference in ERN was noted at the FCz electrode. C) The N200 was significantly more negative in OCD groups than HC at P2 and, D) no between-group difference in N200 was noted at the FCz electrode. E) The P300 was significantly more positive in OCD groups when compared to HC at F3 and, E) no between-group difference in P300 was noted at Pz. (ERN – error related negativity, OCD – obsessive compulsive disorder, HC – healthy control)

### References

[1] G. Hajcak, N. McDonald, and R. F. Simons, "Anxiety and error-related brain activity," *Biological psychology,* vol. 64, no. 1-2, pp. 77-90, 2003.

[2] J. Towey *et al.*, "Event-related potential and clinical correlates of neurodysfunction in obsessive-compulsive disorder," *Psychiatry Research,* vol. 49, no. 2, pp. 167-181, 1993.

[3] C. Andreou *et al.*, "P300 in obsessive–compulsive disorder: source localization and the effects of treatment," *Journal of psychiatric research,* vol. 47, no. 12, pp. 1975-1983, 2013.
